# Revealing Cellular Heterogeneity and *In Vitro* Differentiation Trajectory of Cultured Human Endometrial Mesenchymal-like Stem Cells Using Single-cell RNA Sequencing

**DOI:** 10.1101/2020.04.03.004523

**Authors:** Dandan Cao, Rachel W.S. Chan, Ernest H.Y. Ng, Kristina Gemzell-Danielsson, William S.B. Yeung

## Abstract

Endometrial mesenchymal-like stem cells (eMSCs) are adult stem cells contributing to endometrial regeneration. One set of perivascular markers (CD140b^+^CD146^+^) have been widely used to enrich eMSCs. Although eMSCs are easily accessible for regenerative medicine and have long been studied, their cellular heterogeneity and molecular program controlling their expansion and differentiation in vitro remains largely unclear. In this study, we applied 10X genomics single-cell RNA sequencing to eMSCs cultured in vitro after microbeading from 7 donors to investigate cellular heterogeneity in an unbiased manner. Corresponding clonogenic progenies of eMSCs after culture for 14 days were also sequenced to construct the in vitro differentiation trajectory of eMSCs. Transcriptomic expression based clustering revealed several subpopulations in eMSCs. Each subpopulation manifested distinct functional characteristics associated with immunomodulation, proliferation, extracellular matrix organization and cell differentiation. Pseudotime trajectory analysis on eMSCs and their differentiated progenies identified *in vitro* differentiation hierarchy of eMSCs. Further ligand-receptor pair analysis found that WNT signaling, NOTCH signaling, TGF-beta signaling and FGF signaling were important regulatory pathways for eMSC self-renewal and differentiation. By comparing eMSCs to Wharton’s Jelly MSCs and adipose-derived MSCs, we found these 3 kinds of MSCs expressed largely overlapping differentiation (CD) genes and highly variable genes. In summary, we reveal for the first time high molecular and cellular heterogeneity in cultured eMSCs, and identify the key signaling pathways that may be important for eMSC differentiation.

## INTRODUCTION

The uterine cavity is lined by the endometrium, which is shed off and regenerates in each menstrual cycle. This remarkable physiological remodeling occurs about 400 times in a woman’s reproductive life. Adult stem cells are undifferentiated cells found throughout the body after development. They proliferate and differentiate to replenish dying cells and to regenerate damaged tissues. In endometrium, stromal stem cells was firstly identified as clonogenic cells with multiple lineage differentiation potential [1]. Endometrial stromal stem cells exhibited properties similar to that of mesenchymal stem cells (MSCs) in other tissues in terms of clonogenicity, fibroblast-like morphology, surface markers phenotype and multipotency. Thus they are called endometrium mesenchymal-like stem cells (eMSCs).

MSCs, including eMSCs, exhibit great differentiation potential and immunomodulation ability enabling them for cell therapeutic use. Among MSCs, eMSCs are the only one that can easily be obtained from women each month without use of analgesics. A woman can use her own eMSCs for therapy when needed. Therefore, eMSCs have been tested as an alternative source for cell therapies. Transplantation of human eMSCs to mouse and primate models of Parkinson’s disease significantly increases the dopamine level when compared to sham transplanted controls [2, 3]. In addition, eMSCs have been also studied for regenerative medicine in other diseases including diabetes, cardiac diseases and cartilage injury [4].

Lessons from clinical trials of MSCs show that differences in preparation of MSCs such as culture and expansion method affect treatment efficacy of MSCs [5]. For instance, bone marrow derived MSCs exhibit cellular heterogeneity during expansion *in vitro* [6], and MSCs from different clones exhibited substantial variation in differentiation potential [7]. Previous studies on eMSC assumed that eMSCs were a homogenous population. Our recent study shows the presence of cellular heterogeneity in eMSCs. Apart from inter-donor variation in clonogenic ability of eMSCs, a higher proliferative ability is found in eMSCs from endometrium at the menstrual phase than those from other phases of the cycle, suggesting the presence of a quiescent subpopulation and an activated subpopulation [8]. Thus, knowledge on the cellular heterogeneity of in vitro expanded eMSCs is urgently needed for standardization before applying the cells in clinical therapy.

The recent development of single-cell RNA sequencing (scRNA-seq), which combines single-cell isolation techniques with RNA-seq, creates an opportunity to study the transcriptomes of individual cells enabling clear distinctions between subpopulations, and thorough assessment of gene transcripts in an unbiased manner [9]. In this study, we aim to characterize the gene expression in the CD140b^+^CD146^+^ eMSC population at single cell resolution. We identified several subpopulations of cultured eMSCs and characterized the molecular programs in these subpopulations. *In vitro* differentiation trajectory analysis revealed the developmental states as well as key signaling pathways controlling the differentiation process from eMSCs to differentiated progenies. Our study for the first time fills the knowledge gap on understanding the cellular heterogeneity of eMSCs and provides guidance for future standardization of eMSCs in *in vitro* expansion.

## RESULTS

### Overview of single-cell transcriptomic data

Endometrial aspirates from the menstrual phase of three women and four full-thickness endometrium from the secretory phase of four women were used in this study (**Table S1**). After enzyme dispersion of the tissues, the CD140b^+^CD146^+^ cells were obtained from endometrial stromal cells by serial magnetic microbeading [8, 10]. They were subjected to scRNA-seq on a 10X genomics platform. In addition, single cell-derived colonies from the eMSCs of one secretory sample were also subjected to scRNA-seq (**Table S1**). ScRNA-seq libraries for each sample were constructed independently (**Table S1**).

After quality control of the raw sequencing tags, we obtained around 5×10^8^ sequence reads for each sample (**Figure S1B, Table S2**), with 54.5-72.9% confidently and uniquely mapped to the human reference transcriptome GRCh38 (NCBI) (**Table S2**). In total, 29,438 cells were detected from the 7 samples with 1.29×10^5^ mean reads, 2.9×10^4^ mean unique molecular identifiers (UMIs) and 4,548 mean genes per cell (**Figure S1B, Table S2**). The variation observed in cell number might be due to differences in cell number loading and use of different reagent versions for single cell encapsulation and library preparation for samples (**Figure S1B, Table S2**). Despite differences in the number of cells per sample, there was little variation in the total number of genes detected per sample (**Figure S1B, Table S2**). These results suggested that the sequence depth in our data achieved approximately maximum total gene detection in the samples.

### Variation dominated by cell cycle and donor effect in eMSCs

Based on the sequencing data, 7 eMSC samples were firstly aggregated with depth normalization and chemistry batch correction using the *cellranger* aggr pipeline. Median absolute deviation (MAD) quality control matrices were then applied to filter low quality cells and genes (**Table S3**). Finally, we got 20,646 cells expressing 13,406 genes from the 7 eMSC samples for subsequent analysis (**Table S3**). Typical marker expression were checked in this cell population and the results showed high expression of stromal cell markers *VIM* and *S100A4* in cells from each sample, absence of endothelial cell marker *CD34,* and little expression of epithelial cell marker *EPCAM*, immune cell markers *PTPRC* (CD45) and *CD14* (**Figure 1A**), confirming high purity of the isolated stromal cells with minimal contamination by other cells. Further inspection showed high expression of MSC markers *ITGB1* (CD29), *CD44*, *NT5E* (CD73), *THY1* (CD90) and *ENG* (CD105) in each sample validating the MSC identity of the studied population (**Figure 1A**). Expression of these markers varied across the samples indicating donor heterogeneity (**Figure 1A**).

**Figure 1.**
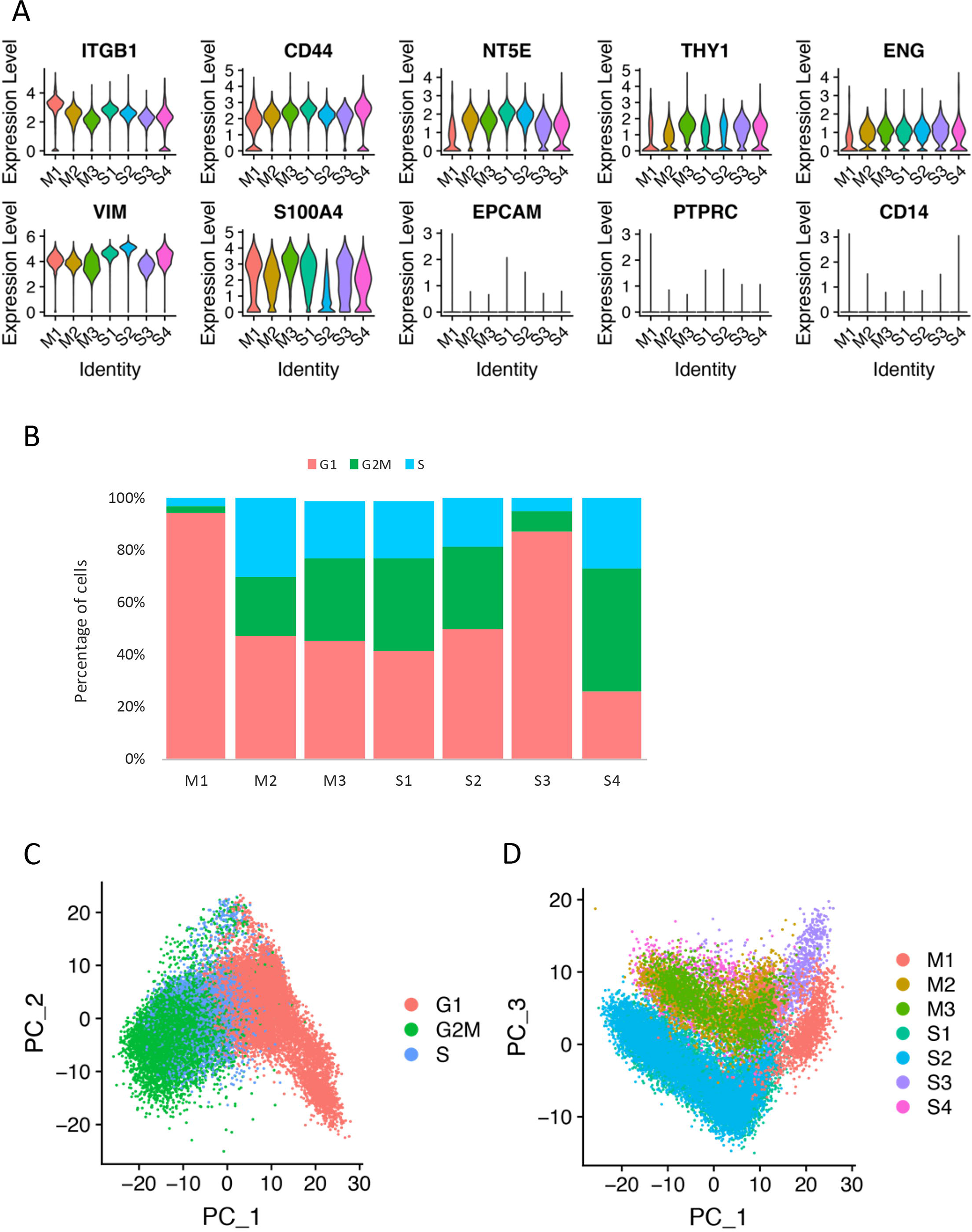
Overview of single-cell RNA sequencing. (A) Violin plots for expression of MSC markers, stromal cell markers, epithelial cell marker, immune cell markers in 7 eMSC samples. (B) Phases of cell cycle distribution in each sample. (C) PCA plot on the first two principal components showing the separation of cells by cell cycle phase. (D) PCA plot on the first and third principal components showing the separation of cells by donor/batch.

To further identify the major variance in the eMSC population at single cell level, we performed cell cycle classification and principal component analysis (PCA). Cell cycle phases (G1, G2M, S) were assigned to each cell using the Seurat package [11, 12]. The cell cycle phase distribution in each sample is shown in **Figure 1B**. At sample level, one menstrual phase sample (M1) and one secretory phase sample (S3) manifested quiescent/senescence characteristic with a high proportion of cells at the G1 phase, whereas the remaining samples demonstrated a high proportion of cells in the proliferative phase (**Figure 1B**). From overview of all single cells, PCA results based on normalized gene expression matrix showed that variance of the first 3 principal components were mainly caused by cell cycle effect (**Figure 1C**) and batch/donor effect (**Figure 1D**), which were consistently observed in other single cell studies on MSCs [13–15]. The results indicated that unwanted major source of variation should be removed before investigation on the molecular heterogeneity of eMSCs.

### Identification and biological classification of distinct subpopulations in eMSCs

To remove the cell cycle effect, we scaled the data and regressed out the cell cycle effect by linear regression method implemented in the Seurat package [11, 12]. The fastMNN [16] was used to remove the batch effect. After regression and correction, nonlinear dimensional reduction by Uniform Manifold Approximation and Projection (UMAP) of the data showed that both the cell cycle and the batch/donor effect were largely mitigated (**Figure S2A, S2B**).

After removal of the unwanted source of variation, candidate population clustering by a shared nearest neighbor (SNN) graph-based approach revealed 8 subpopulations (SP1-SP8) for all the 20,646 cells (**Figure S2C**). Cell distribution in each subpopulation is shown in **Table S4**. Quality control post-clustering showed that cells in SP7 were mostly from sample S2, cell number in SP8 was less than 100, and UMI count (median = 877) and gene number (median = 595) of SP6 were extremely low when compared to the remaining subpopulations, suggesting unreliability of these three subpopulations (**Figure S2C, Table S4**). Further analyses were made on the remaining 5 high quality candidate subpopulations SP1-SP5 (**Figure 2A, Table S4**). The composition of SPs for each sample was shown in **Figure S2D**. No menstrual phase-specific subpopulations were present. Pearson’s correlation analysis on average gene expression of the subpopulations showed that SP3 was a distinct subpopulation, while SP1 and SP4 were highly correlated (**Figure 2B**). The markers of MSC, stromal cells and pericytes were expressed at a relatively comparable level in all candidate subpopulations except SP2 which exhibited a relatively low expression of *THY1* (CD90) and *S100A4* (**Figure S3A**). A comprehensive analysis on individual cells was performed to assess the self-renewal, multiple lineage differentiation and immunomodulation ability for each subpopulation by calculating the mean expression of markers in each category as defined in **Table S5,** which were retrieved from STEMCELL™ TECHNOLOGIES website (https://stemcell.shinyapps.io/qpcr_tool/) and an online report [13]. The results demonstrated that all subpopulations expressed a high level of MSC markers. SP1 and SP3 expressed stemness markers (**Figure S3B**). Moreover, all subpopulations possessed a high potential to differentiate into chondrogenic, osteogenic and neurogenic lineages, but limited adipogenic lineage differentiation capacity (**Figure S3B**). Expression of immunosuppression markers was highest in SP5 and minimal in SP1 and SP4 (**Figure S3B, Table S5**).

**Figure 2.**
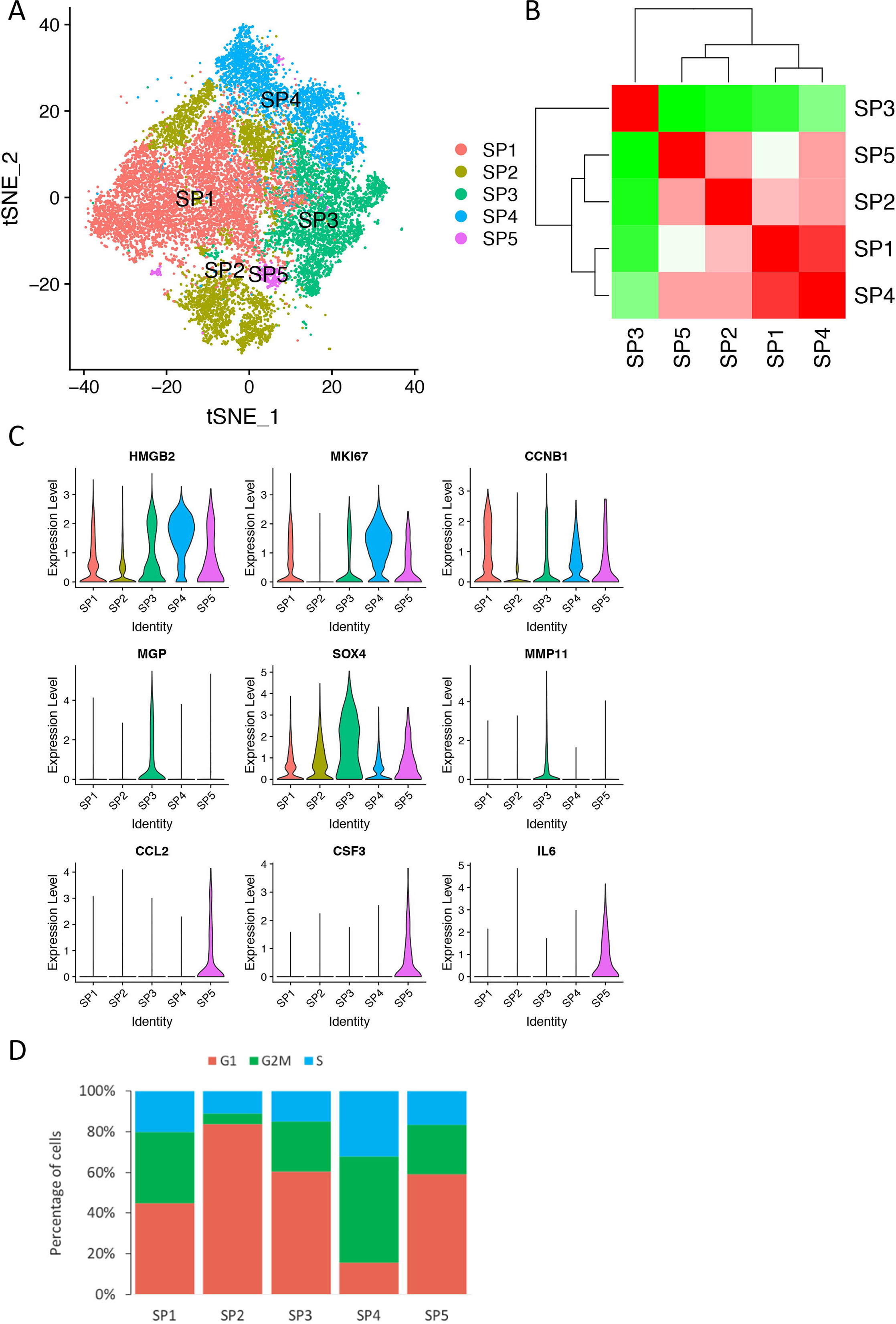
Identification of subpopulations in eMSCs. (A) A t-SNE plot for 20,646 cells colored by subpopulation assignment. In total, 5 subpopulations were identified. (B) Pearson’s correlation between the average gene expression profiles of 5 subpopulations. (C) Violin plots for the expression of top differentially expressed genes (DEGs) in different subpopulations including HMGB2 and MKI67 in SP4, CCNB1 in SP1, MGP, SOX4, and MMP11 in SP3, and CCL2, CSF3 and IL6 in SP5. (D) Phases of cell cycle distribution in each subpopulations.

To further investigate the biological processes underlying the transcriptional classification of the cell subpopulations, we identified the upregulated genes (UGs) in each subpopulation by comparing it to the remaining clusters with the Wilcoxon Rank Sum test. The UGs with an average log2 fold-change greater than 0.25 and a Bonferroni-corrected P-value threshold (P < 3.73×10^−6^) were subjected to downstream analysis. In total, we identified 46, 100, 206, 183 and 100 UGs for SP1, SP2, SP3, SP4 and SP5, respectively (**Table S6**). Interestingly, typical proliferative markers *HMGB2* and *MKI67* were highly expressed in SP4 (**Figure 2C**), implying a high proliferative potential of this population. Consistently, cell cycle analysis showed a low percentage of G1 phase cells (16%) in SP4 (**Figure 2D**). SP1, a population correlated to SP4, also expressed high level of *CCNB1*, a gene essential for control of cell cycle at the G2/M (mitosis) transition (**Figure 2C**). In silico gene ontology (GO) enrichment analysis of UGs in SP1 and SP4 identified pathways associated with regulation of mitosis/cell cycle and DNA replication (**Figure 3A, 3D, Table S7**). Additionally, SP1 contained less G1 cells when compared to SP2, SP3 and SP5 (**Figure 2D**).

**Figure 3.**
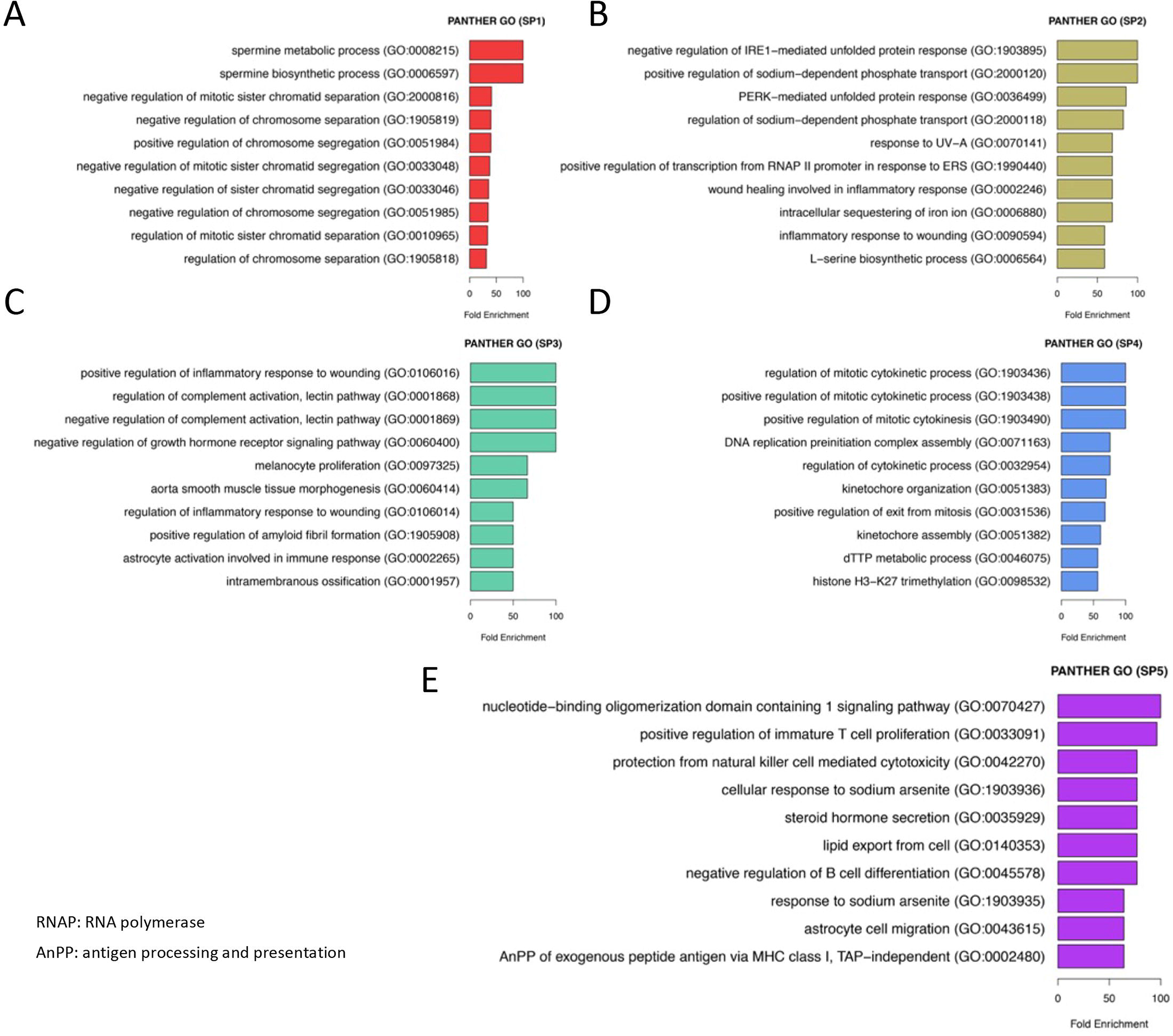
Gene ontology analysis of differentially expressed genes in each subpopulation. (A-E) Top significant enriched biological processes from PANTHER gene ontology analysis based on UGs identified in (A) SP1, (B) SP2, (C) SP3, (D) SP4, and (E) SP5.

Unlike SP1 and SP4, SP3 and SP5 showed medium proliferation ability (**Figure 2C, 2D**) but high immunomodulation capacity. Among the UGs in SP5, the top ones were all inflammatory cytokines such as *CCL2*, *CSF3*, *IL6*, and different *CXCLs* (**Figure 2C, Table S6**). GO analysis on the UGs revealed that the SP5 cells manifested biological functions relating to immunomodulation like regulation of T cell proliferation and B cell differentiation (**Figure 3E, Table S7**). We speculated that the SP5 cells could interact with the immune system, thus could be a promising population for therapeutic use. GO analysis of SP3 also showed superior immunomodulation activities including regulation of inflammatory responses and complement activation (**Figure 3C, Table S7**). We noted that SP3 highly expressed the gene *MGP* (**Figure 2C**), an inhibitor of bone morphogenic proteins (BMPs) signaling known to immunomodulate functions of a subpopulation of mouse MSCs [17]. Intriguingly, the UGs of SP3 are enriched in biological processes including tissue morphogenesis, extracellular matrix (ECM) organization, and cell differentiation (**Figure 3C, Table S7**). SP2 possessed the highest proportion of G1 cells (**Figure 2D**), expressed genes related to responses to cellular unfolded protein or endoplasmic reticulum stress (ERS) and underwent activities involving cellular response to glucose starvation and oxidative stress (**Figure 3B, Table S7**), implying discounted proliferation ability and high mitochondrial activity in this subpopulation.

Next the linkage of the identified subpopulations with primary endometrial cell lineage was investigated. We collected and analyzed single cell data from a primary endometrial sample. Unsupervised clustering analysis revealed stromal cells, epithelial cells, endothelial cells, macrophages and other subpopulations in the primary endometrium (**Figure S4A**). MetaNeighbor analysis linked SP1 and SP4 to the epithelial cell lineage, SP2 and SP3 to the stromal cell lineage, and SP5 to the macrophages (**Figure S4B**). The results not only supported the superior immunomodulation ability of SP5, but also revealed the differentiation potential of SP2 and SP3 into stromal cells, and the trans-differentiation potential of SP1 and SP4 into epithelial cells.

In summary, we identified 5 subpopulations in cultured eMSCs unbiasedly, and highlighted their unique properties in terms of proliferation, immunomodulation, mitochondrial activity, cell differentiation and extracellular matrix organization.

### Construction of *in vitro* differentiation trajectory for eMSC

Similar to MSCs, eMSCs also show a finite proliferation period followed by senescence or differentiation. We studied the differentiation of eMSCs by culturing the sample S3 for an additional 14-day and performed scRNA-seq on their clonogenic progenies (S3C) (**Table S1, Figure S1B**). In this paired samples, the expressions of all MSC markers (*ITGB1*, *CD44*, *NT5E*, *THY1*, *ENG*) in cells from S3C were lower than that from S3 (**Figure S5A**). Notably, the eMSC isolation markers (CD140b, CD146) used in the study were dramatically decreased in the S3C cells when compared to their parental S3 cells (**Figure S5A**), suggesting the loss of eMSC identity in S3C.

After quality control processing as described above (**Table S8**), 3,928 cells and 13,967 genes were obtained from the 2 samples for analysis. The number of genes and the number of total UMIs per cell were comparable between the two samples (**Figure S5B**). Cell cycle phase distribution in each sample was shown in **Figure S5C**. Similar cell cycle effect and donor effect existed in the samples (**Figure S5D, S5E**), and the same strategy was applied to remove the unwanted source of variation (**Figure S5F**). There was little overlap between the S3 and the S3C cells because majority of the S3C cells had lost their expression of eMSC markers (**Figure S5G**).

Clustering of the corrected dataset revealed 7 primary clusters (**Figure 4A, Table S9**). Post-clustering quality control abandoned 2 clusters with low gene numbers and UMI counts (**Figure S6A**). The Pearson’s correlation on average gene expression in the remaining 5 clusters is shown in **Figure 4B**. We compared the clusters from S3 and S3C to the subpopulations of the 7 eMSCs samples described above using MetaNeighbor [18]. The results revealed that cluster 3 was similar to SP3 with an AUROC score 0.85, while cluster 4 was similar to SP2 with an AUROC score 0.86. Marker expression analysis showed that cluster 0, 2, 4 expressed lower levels of MSC markers *ITGB1*, *NT5E* and *ENG*, and pericyte markers *PDGFRB*, *MCAM* and *ACTA2* when compared to cluster 1 and 3 (**Figure S6B**). To construct the *in vitro* lineage differentiation of eMSCs, we performed cell trajectory and pseudotime inference analysis on clusters 0-4 using the Monocle 2 [19]. The results revealed that majority of the cells from cluster 3 and a proportion of cells from cluster 1 were located at the root, while majority of the cells from cluster 0 and 2 were distinctly at the terminations representing more differentiated cells (**Figure 4C-4E**). Cells along the trajectory were mostly from cluster 1 and 4, indicating their roles as intermediates (**Figure 4C-4E**).

**Figure 4.**
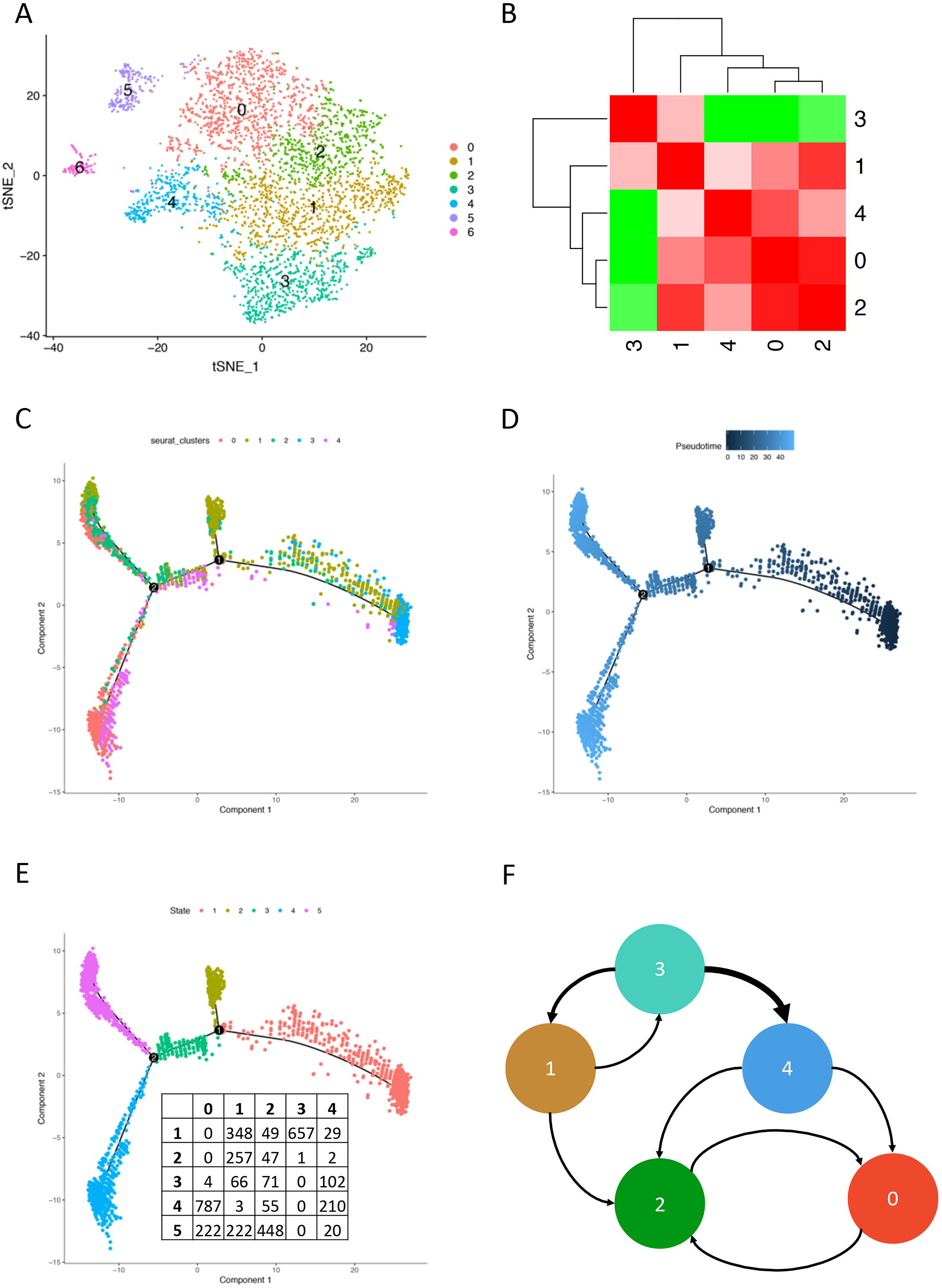
Clustering and trajectory analysis of single cells from eMSC sample S3 and its paired colonogenic sample S3C. (A) A t-SNE plot for 3,928 cells colored by cluster assignment. In total, 7 primary clusters were identified. (B) Pearson’s correlation between the average gene expression profiles of 5 clusters of high quality. (C-E) Pseudotime analysis of single cells using Monocle 2 identified cells on the tree colored by (C) cluster assignment, (D) pseudotime and (E) state. HVGs used for clustering were used to construct the pseudotime tree. The cells on the right side (dark blue) of the pseudotime tree are less differentiated, while those on the left side (light blue) are more differentiated. Overlaying cluster information (C) shows cells from cluster 3 are eMSCs and cells from cluster 0 and 2 are differentiated eMSCs. Cell states (E) are cells on the same branch with similar pseudotime values. Cell distribution across clusters (columns) in each state (rows) were listed. (F) Transitional potential between different clusters identified from scGPS analysis. The weight of the arrows is relative to the percentage summarized in Table S10 (thicker indicates higher percentage).

To quantitatively estimate the proportion of cells with potential to transit from one cluster to another, we utilized scGPS (nbootstrap = 100) (https://github.com/IMB-Computational-Genomics-Lab/scGPS) to perform the transitional analysis. The result showed that cells in cluster 3 possessed great potential to progress to cells in cluster 4 and 1; cluster 4 had the potential to transit to cluster 0 and 2; and cluster 1 could also transit to cluster 2 (**Figure 4F, Table S10**). In addition, possibilities of inter-conversion of cells between cluster 3 and 1 and between cluster 0 and 2 were noted (**Figure 4F, Table S10**).

The findings from Pearson’s correlation, pseudotime trajectory inference and scGPS transitional potential evaluation indicated that eMSCs in cluster 3 developed into differentiated eMSCs in cluster 0 and 2 through intermediates in cluster 1 and cluster 4 during *in vitro* differentiation,

### Molecular signatures involved in *in vitro* differentiation of eMSC

To systematically study interactions between different subpopulations identified during *in vitro* differentiation, we applied the CellPhoneDB [20] to identify significant subpopulation-specific ligand-receptor pairs (**Table S11**) and found wide existence of ligand-receptor interactions involving FGF signaling, WNT-signaling, NOTCH signaling and TGF-beta signaling among the cell clusters (**Figure 5A, Table S11**). For example, *WNT5A* was highly expressed in cluster 3 and 1, and could interact with the other clusters through *PTPRK*, *FZD2* and *FZD6* (**Figure 5A**). In addition, cluster 3 and 1 also expressed *NOTCH3*, which could interact with *DLL3* expressed in cluster 0-2. *TGFB1* was expressed highly in clusters 0 and 2 and could act through its receptor TGFB receptor 1 and 2 expressed in other clusters (**Figure 5A**). Interestingly, *TGFB1* induces differentiation of human bone marrow-derived MSCs [21]. We further checked the dynamic expression of molecules of WNT and NOTCH signalings during pseudotime development. Decreasing expression of *FN1*, *NOTCH2*, *NOTCH3*, *WNT5A*, *FZD3*, *FZD4*, and the increasing expression of *DLL3* and *TGFB1* were observed with differentiation (**Figure 5B**), supporting their involvement in eMSC differentiation. More interaction pairs are summarized in **Table S11**.

**Figure 5.**
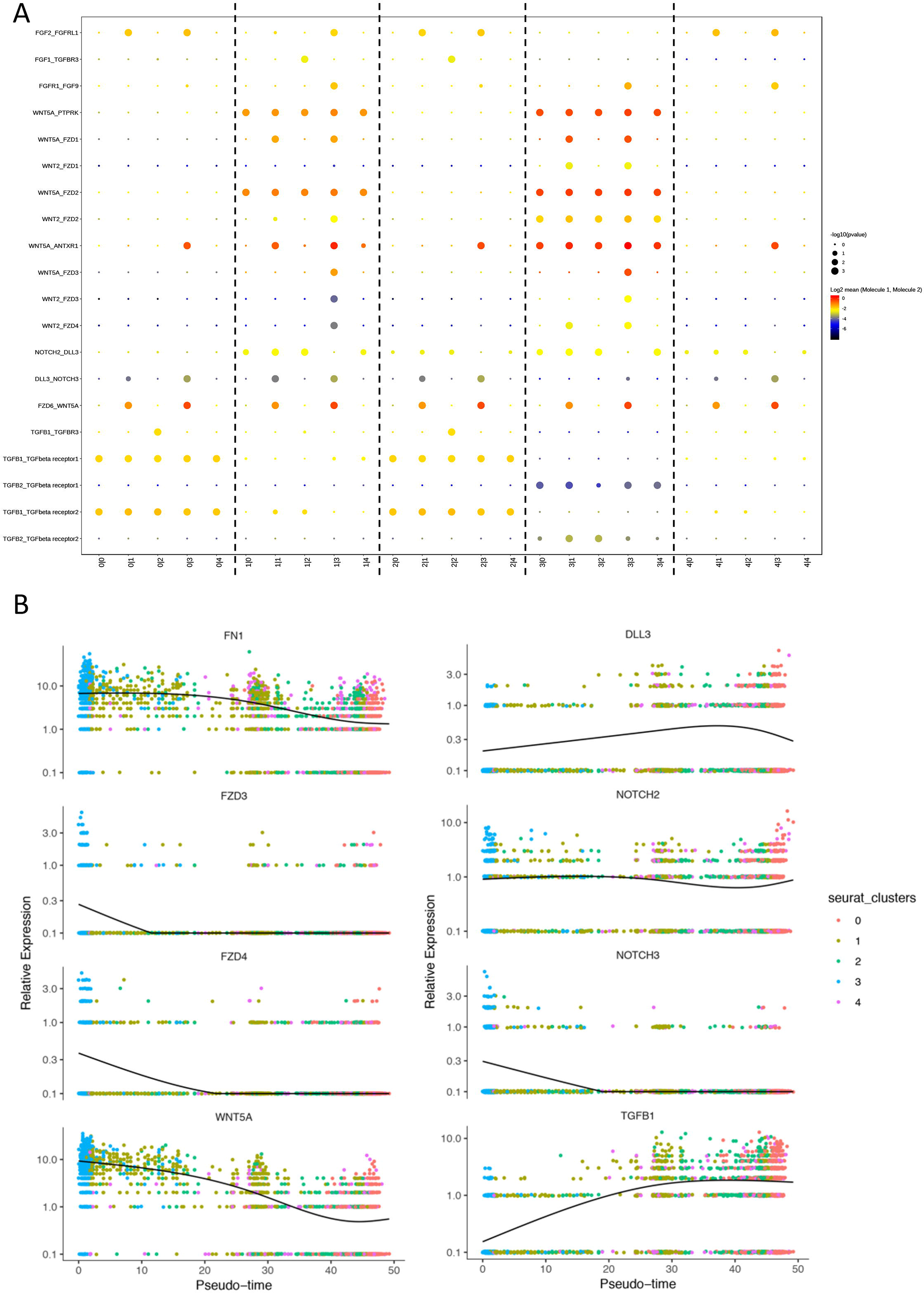
Multiple regulatory pathways identified during *in vitro* eMSC differentiation. (A) Overview of selected ligand-receptor interactions among clusters representing eMSCs and differentiated eMSCs. P values indicated by circle size, scale on right. The means of the average expression level of interacting molecules are indicated by color. In the interaction pair, the former molecule expressed in the first subpopulation while the latter molecule expressed in the second subpopulation (x-axis, e.g 0|1). (B) Dynamic expression of genes from different signaling pathways during pseudotime.

### Transcriptomic comparison of eMSCs to MSCs of other sources

Endometrium is an easy-to-access source for MSCs. Previous studies demonstrated heterogeneity of MSCs from other sources [13, 15, 22, 23]. Here for the first time, we compared the transcriptomes among eMSCs, adipose-derived MSCs (ADMSCs), and umbilical cord-derived (Wharton’s Jelly) MSCs (WJMSCs) at single cell level.

We firstly identified the top 50 cluster of differentiation (CD) genes (ranked by average expression) expressed in the whole population of 7 eMSCs samples (**Figure 6A**). Although *ITGB1*, *CD44*, *NT5E*, *THY1* and *ENG* were widely accepted to be used as markers for MSC, their expression were not the highest in eMSCs (**Figure 6A**). Thirty-seven of the CD genes shared with the top 50 CD genes for ADMSCs and WJMSCs retrieved from a pre-published study [13] (**Figure 6B, Table S12**). The correlation coefficient is 0.72 between eMSCs and ADMSCs, 0.57 between eMSCs and WJMSCs, and 0.44 between ADMSCs and WJMSCs according to the Spearman’s ranking correlation analysis based on the top 50 CD genes. We used Seurat to identify the highly variable genes (HVG), which represent the molecular heterogeneity within a studied cell population. We could see that majority of the HVGs (544/770) of WJMSCs were present in HGVs of eMSCs (**Figure 6C**). High overlap of HVGs from WJMSCs and ADMSCs have been reported [13]. The overlapping HVGs of different MSCs indicated their critical roles in functioning of MSCs; meanwhile, the unshared HGVs might serve as regulators which mark the differences between different types of MSCs.

**Figure 6.**
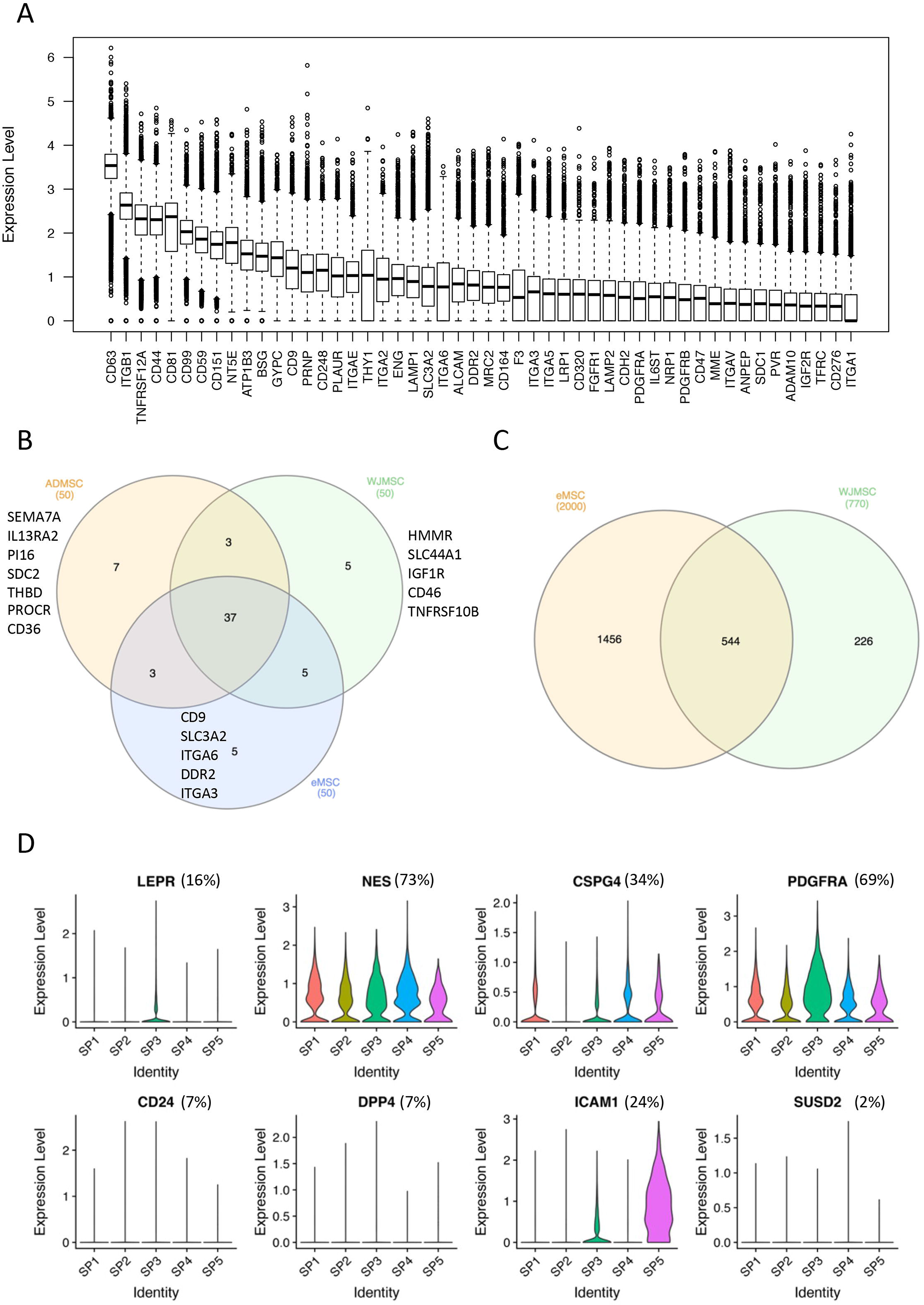
Comparing eMSCs to reported MSCs at single-cell level. (A) Boxplot showing the top 50 CD genes ranked by average normalized expression in the whole eMSCs population. (B) Venn diagram showing the overlap relationship among top 50 CD genes for cultured eMSCs, ADMSCs and WJMSCs. (C) Venn diagram showing the top 2000 highly variable genes (HVGs) in eMSCs overlap with published HVGs identified for WJMSC. Majority of HVGs in WJMSC were present in that of eMSCs. (D) Violin plots showing expression of reported MSC subpopulation markers in subpopulations identified in eMSCs. Percentage represented the ratio of total positive cells expressing that marker with UMI greater than 0 among total cells of eMSCs.

Several surface markers have been used to isolate subpopulations of MSCs. For example, *LEPR*, *NES*, and *CSPG4* (*NG2*) are marker for the subtypes of bone marrow MSCs (BMSCs) [24–27] while *PDGFRA*, *CD24*, *DPP4*, and *ICAM1* are markers for subpopulations of ADMSCs [23, 28, 29]. These markers had different expression patterns in eMSC subpopulations (SP1-SP5) identified from our 7 eMSC samples (**Figure 6D, Table S13**). High percentage of eMSCs expressed BMSC subtype marker *NES* (73%) and ADMSC subtype marker *PDGFRA* (69%) (**Figure 6D, Table S13**). Although less (34%) eMSCs expressed the BMSC subtype marker *CSPG4*, it is expressed in all eMSC subpopulations except SP2 (**Figure 6D, Table S13**). ADMSC subtype markers *CD24*, *DPP4* were rarely expressed in eMSCs (**Figure 6D, Table S13**). Interestingly, a BMSC subtype marker *LEPR* was highly expressed only in SP3, while an adipose MSC subtype marker *ICAM1* were expressed only in SP3 and SP5 (**Figure 6D, Table S13**). The expression of another eMSC marker *SUSD2* was negligible in our eMSCs, suggesting little overlap between CD140b^+^CD146^+^ eMSCs and SUSD2^+^ eMSCs (**Figure 6D, Table S13**) consistent with a previous report [30]. The functional implications of different expression patterns of MSC subtype markers in CD140b^+^CD146^+^ eMSCs subpopulations remain to be determined.

## DISCUSSION

Endometrial MSCs play critical roles in the cyclic regeneration of human endometrium [31]. Two sets of markers can enrich for eMSCs [30, 32], and FACS analysis showed little overlap between these two eMSC populations [30], suggesting heterogeneity in the whole eMSC population. Heterogeneity was also observed within cultured eMSCs in terms of proliferative potential [32, 33]. Although the heterogeneity of eMSCs is widely recognized, little has been done to identify and characterize their subpopulations. To address this, we generated and analyzed a large single-cell transcriptomic dataset from cultured CD140b^+^CD146^+^ eMSCs of 7 donors. Additionally, scRNA-seq data of differentiated eMSC progenies were also analyzed to study the *in vitro* differentiation trajectory of eMSCs.

Utilizing high quality single cell data from 20,646 cells collectively expressing 13,406 genes, we revealed that the major sources of variation in our eMSC population were from cell cycle effect and donor/batch effect, consistent with other studies on MSCs. Cell cycle phase distribution differences were observed among different donors. Whether the differences were inherent across different individuals or caused by *in vitro* culture requires further investigation.

We identified 5 subpopulations (SP1-SP5) of CD140b^+^CD146^+^ eMSCs with different potential in proliferation, immunomodulation, ECM organization and mitochondrial activity, and differentiation. An outstanding population, SP5, had superior immunomodulation abilities expressing a lot of cytokines including *CXCL1*, *CXCL3*, *CXCL6*, *CXCL8*, *CCL2*, *CSF2*, *CSF3*, *IL6*, *IL1B*, related to immunosuppression and angiogenesis [34–37]. Therefore, this subpopulation could be a promising cell source for therapeutic use.

The subpopulation SP3 also showed immunomodulation activities such as complement activation. Interestingly, SP3 highly expressed *MGP*, which was also highly expressed in a subpopulation of mouse bone marrow MSCs [38]. *MGP* is an immunomodulator suppressing activated T cells *in vitro* [17]. The function of *MGP* in eMSCs required further investigation.

In addition to immunomodulation, SP3 was also related to other functions including ECM organization, tissue morphogenesis, and development. SP3 had a high stemness score and the potential to differentiate into stromal cells. When compared to the remaining subpopulations, SP3 contained most primary eMSC-specific genes identified in *Barragan’s* (data not shown) work [33]. Taken together, SP3 is a subpopulation possessing primary eMSC properties, and is the most undifferentiated cell subpopulation among the cultured CD140b^+^CD146^+^ eMSCs. However, whether SP3 exists in primary endometrium requires further validation.

SP1 and SP4 are two related subpopulations with high proliferative capacity. They were identified as two independent subpopulations because SP4 expressed higher levels of proliferative markers (*MKI67*, *TOP2A*, *HMGB2*) than SP1. Intriguingly, the SP4 cells had the highest proliferation capacity and were enriched in the menstrual phase samples (*chi-squared test*, p < 0.0001). We have demonstrated that eMSCs from the menstrual phase have a higher proliferation ability than those from the secretory phase [8], and may be attributed to the enrichment of the SP4 cells in the menstrual phase. In addition, SP1 and SP4 showed a trans-differentiation potential to the epithelial cell lineage (**Figure S4B**).

SP2 showed limited proliferation but were associated with high unfolded protein responses and high metabolic activities. Relatively higher proportion of mitochondrial reads was observed in SP2 than in other subpopulations (**Figure S2C, Table S4**). High mitochondrial activities are observed in cancer cells [39]. Additionally, mitochondrial metabolism is a key regulatory mechanism in stem cell fate decision [40]. *Samual* and coworkers reported that mitochondrial metabolism is higher in endovascular progenitor cells than that in differentiated endothelial cells [41]. On the other hand, higher mitochondrial reads in scRNA-seq might be due to presence of cells of low quality resulting from damages during cell dispersion. MetaNeighbor analysis showed that SP2 was correlated with the differentiated cluster along the differentiation trajectory, suggesting that SP2 was likely to be a subpopulation undergoing senescence/dying rather than a progenitor subpopulation. However, further investigation on SP2 is still required to distinguish these two possibilities.

Cell distribution across SPs (**Figure S2D, Table S4**) showed that no menstrual cycle stage-specific subpopulation was identified. However, different proportion of SPs in each sample was observed in this study (**Figure S2D**). Specifically, menstrual phase samples M1, M2 and secretory sample S2 contained more SP4 cells, while menstrual sample M1 and secretory sample S3 contained more SP3 cells when compared to the remaining samples. This observation suggested that our samples individually undergo clonal selection and each sample in our study might be on different stages of clonal selection, a phenomenon reported in MSCs in a previous study [42].

To obtain large amount MSCs for clinical use requires *in vitro* expansion because of limited supply of primary MSCs. However, *in vitro* expansion has large inter-sample variations and causes functional loss of the cells [43–46]. Thus, understanding of *in vitro* differentiation is critical. In this study, we investigated the *in vitro* differentiation of eMSCs at single-cell level. Through clustering and trajectory analysis, we for the first time demonstrated the differentiation path of eMSCs: different clusters of cells were differentiated from the parental eMSCs through less differentiated intermediates. Further ligand-receptor analysis identified important signaling pathways regulating the differentiation. For example, the WNT signaling pathway was identified in the undifferentiated/less differentiated clusters. Recently, we demonstrate that *WNT5A* is a niche factor that promotes self-renewal of eMSCs through *FZD4* and canonical WNT-signaling [10]. Here other WNT receptors like *FZD3* were identified, highlighting the importance of dissecting the heterogeneity of eMSCs. In addition to WNT signaling pathway, NOTCH signaling, TGF-beta signaling and FGF signaling pathways were also identified. They are all important pathways regulating MSC self-renewal and differentiation [47–49]. The interactive molecules of the pathways among clusters identified here could guide future studies investigating the regulatory mechanisms of eMSC differentiation.

Several studies compared transcriptomes between different types of MSCs [13, 50, 51]. We compared for the first time the mRNA profile of eMSCs to other types of MSCs at single-cell level. MSCs are defined by a combination of surface markers [52]. Thus, we firstly compared the top 50 expressed CD genes among eMSCs, ADMSCs and WJMSCs, since they are all cultured primary MSCs and processed in a 10X genomics platform. Although a majority of the top expressed CD genes were shared among them, the expression ranking was different. Interestingly, the typical CD genes that define MSCs were not expressed highest in all the three groups, indicating that a new marker set for identifying MSCs could be considered. Similarities and differences in HVGs were also observed among different types of MSCs. We found that the eMSC subpopulations relating to proliferation and immunomodulation were also found in the WJMSCs [13]. However, the subpopulation with high mitochondrial activity (SP2) was unique to eMSCs.

In the past years, MSCs marked by specific markers were studied. Through investigation on the expression of bone marrow MSC subtype markers *LEPR*, *NES*, *CSPG4* (NG2) [24–27], adipose MSC subtype markers *PDGFRA*, *CD24*, *DPP4*, *ICAM1* [23, 28, 29] and eMSC subtype marker *SUSD2* [30], we found that these markers exhibited different expression patterns among the CD140b^+^CD146^+^ subpopulations. Some were widely expressed in each subpopulation (*PDGFRA*, *NES*, and *CSPG4*); some were expressed at a minimal level (*CD24*, *DPP4*, and *SUSD2*); while some of them expressed only in distinct subpopulations (*LERP* and *ICAM1*). The results further indicated the differences and similarities among different types of MSCs.

Here we report the first single cell sequencing study on eMSCs based on a large offset of cells. In addition, cells were obtained from several donors at different menstrual phases, ensuring the comprehensiveness. Undoubtedly, this large cell atlas of human cultured CD140b^+^CD146^+^ eMSCs provides an essential resource for a better understanding into the nature of eMSCs and guidance for the production of homogenous eMSCs for cell therapy. In the future, more work should be done to determine the generalizability of the present observations by including more samples in the analyses of eMSC differentiation and expanding to other eMSC population.

## METHODS

### Human tissues

Ethical approval was obtained from the Institutional Review Board of The University of Hong Kong/Hospital Authority Hong Kong West Cluster (IRB reference number UW15-128). Each woman signed a written informed consent after fully counselled. Menstrual phase samples were collected by endometrial aspiration from four women with regular menstrual cycles (median age 32; range 31-40 years) attending the infertility clinic on day 2-3 of their menstrual cycle. Full thickness endometrial samples were collected from women with regular menstrual cycles (median age: 50; range: 49-52 years) who underwent total abdominal hysterectomy for benign non-endometrial pathologies. They had not taken hormonal therapy in the past three months before the surgery. The endometrial samples (n = 4) were found to be at the secretory phase of the menstrual cycle assessed by experienced pathologists based on histology of endometrial sections.

### Isolation of endometrial cells

Endometrial tissues were minced into 1 mm^3^ pieces and dissociated in phosphate-buffered saline (PBS) containing collagenase type III (0.3 mg/ml, Worthington Biochemical Corporation, Freehold, NJ, USA) and deoxyribonuclease type I (40 μg/ml, Worthington Biochemical Corporation) in a shaking water bath for 60 minutes at 37°C [53]. After two rounds of digestion, the dispersed cells were filtered through 40μm sieves (BD Bioscience, San Jose, CA, USA), loaded onto Ficoll-Paque (GE Healthcare, Uppsala, Sweden) for removal of red blood cells, cell debris and cell clumps by centrifugation. Anti-CD45 antibody coated Dynabeads (Invitrogen, Waltham, MA, USA) were used to eliminate leukocytes. Stromal cells were negatively selected using microbeads coated with antibody against epithelial cell marker CD368 (EpCAM) (Miltenyi Biotech, Bergisch Gladbach, Germany). Freshly purified stromal cells were plated onto 100 mm dishes coated with fibronectin (1 mg/ml, Invitrogen) containing growth medium containing 10% FBS (ThermoFisher Scientific, Waltham, MA, USA), 1% penicillin (ThermoFisher Scientific) and 1% L-glutamine (ThermoFisher Scientific) in DMEM/F12 (Sigma-Aldrich, St Louis, MA, USA). The stromal cells were expanded in culture for 7–14 days in a humidified carbon dioxide incubator at 37°C. The culture medium was changed every 7 days.

### Magnetic bead selection of endometrial mesenchymal stem-like cells

Isolation of eMSCs (CD140b^+^CD146^+^ cells) was conducted with two separate positive magnetic bead selections [8]. Stromal cells were incubated with Phycoerythrin (PE)-conjugated anti-CD140b antibody at 4°C for 45 minutes. The cells were then incubated with anti-mouse IgG1 magnetic microbeads (Miltenyi Biotech) at 4°C for 15 minutes. The CD140b^+^ cells were collected using the Miltenyi columns with a magnetic field, and cultured for 7 to 10 days in growth medium to allow degradation of the microbeads during cell expansion. The CD140b^+^ cells were then trypsinized and incubated with anti-CD146 microbeads (Miltenyi Biotech) at 4°C for 15 minutes. The CD140b^+^CD146^+^ cells were collected and used for single-cell RNA sequencing and clonogenic culture. For clonogenic assay, 500 CD140b^+^CD146^+^ cells were seeded onto fibronectin coated 10 cm plates and cultured for 14 days.

### Single-cell RNA sequencing

Single-cell RNA sequencing (scRNA-seq) was performed at the Genomics Core, Centre for PanorOmic Sciences (CPOS), The University of Hong Kong. Single cell encapsulation and cDNA libraries were prepared by Chromium™ Single Cell 3’ Reagent Kits v2/v3 and Chromium™ Single Cell A/B Chip Kit. Libraries were sequenced on an Illumina NovaSeq 6000 instrument using paired-end 151 bp.

### Mapping of sequencing reads to human transcriptomes and original cells

High quality sequencing reads for each sample were separately mapped to the human reference genome and transcriptome (GRCh38-3.0.0) using the STAR aligner [54] in the 10X Genomics cellranger pipeline (v3.0.2). Aligned reads were filtered for valid cell barcodes and unique molecular identifier (UMI) during cellranger count process. Gene expression matrix for all samples was generated after between-sample depth normalization using the cellranger aggr.

### Preprocessing of scRNA-seq data

The aggregated single-cell gene expression data was input for the Seurat package [11, 12]. Expression levels for each transcript were determined using the number of UMIs assigned to the transcript. Quality control and filtering steps were performed to remove outlier cells and genes. Cells were discarded if their library size or number of expressed genes, or percentage of mitochondrial reads exceeded the 3x Median Absolute Deviation (MAD). Genes were removed if they were expressed in less than 1% of total cells. In addition, mitochondrial genes and ribosomal genes were also excluded.

### Cell cycle phase classification and cell cycle effect removal

To classify the cell cycle phase of each cell, we firstly assign a score to each cell based on its expression of G2/M and S phase markers [55] using the CellCycleScoring function in Seurat package. Cells were predicated to the G2/M or S phase based on their expression score, while cells expressing neither were likely not cycling and were assigned to the G1 phase. To remove the cell cycle effect, the S scores and G2/M scores were used to regress out cell cycle effect.

### Defining highly variable genes

To define highly variable genes (HVGs), we firstly normalized the data using the Seurat function NormalizeData with method ‘logNormalize’. We then applied the method ‘vst’ of Seurat function FindVariableGenes to identify the top 3000 HVGs for subsequent analysis.

### Dimensionality reduction

Dimensionality reduction was performed after batch correction. We first used the FindVariableFunction of Seurat package to select the top 3000 variable genes. Effects of cell cycle, gene number, total UMIs and percentage of mitochondrial reads were then regressed out when scaling the data. Next, principal component analysis (PCA) was performed to reduce the data to the top 50 PCA components.

### Batch correction

Fast mutual nearest neighbors (fastMNN) [16] correction was performed to remove batch effect among the individuals. Briefly, output PC matrix were input to the fastMNN function implemented in the Seurat package. The data slot ‘mnn’, which contained the corrected matrix were used for downstream clustering analysis.

### Clustering

We conducted a graph-based clustering approach. First, a K-nearest neighbor (KNN) graph was constructed based on the Euclidean distance in mnn space, with refined edge weights between any two cells based on Jaccard similarity using the FindNeighbors function of the Seurat package. Next, the Louvain algorithm was applied to cluster the cells using the FindClusters function of the Seurat package. We visualized the clusters on a 2D map produced with t-distributed stochastic neighbour embedding (t-SNE) or UMAP.

### Differential expression of gene signatures

For each cluster/subpopulation, we used the Wilcoxon Rank-Sum Test to find gene that had significantly different expression when compared to the remaining clusters using the FindAllMarkers function in the Seurat package. Only positive markers were considered. Genes with log fold change larger than 0.25 and Bonferroni correction p values less than 3.73xe-6 were retained and used for further analysis.

### Correlation analysis

We quantified the correlation of single-cell clusters based on average gene expression and ankings of common CD genes among eMSCs, ADMSCs and WJMSCs using the *cor* (method pearson) function in R (3.6.0).

### MetaNeighbor analysis

MetaNeighbor analysis was performed using the the R function MetaNeighbor with default settings [18]. The AUROC (Area under the Receiver Operating Characteristic) scores produced by the MetaNeighbor analysis indicate the degree of correlation between cell groups. An AUROC score of 0.5 means that the probability of correct assignment of a cell’ identity in a binary classification is the same as random guessing.

### Cell trajectory and pseudotime analysis

For pseudotime analysis, we used the Monocle 2 to order cells across the subpopulations based on the HVGs that are used for clustering. The scGPS was used to calculate the percentage of transitional cells between different subpopulations. The top 200 significant differentially expressed genes in each subpopulation identified by the Findmarkers function of the scGPS package were input to scGPS for transitional prediction analysis.

### Cell-cell communication analysis

To systematically analyze the cell-cell communication molecules between different subpopulations in our single-cell data, we performed the CellPhoneDB (a python package) analysis [20]. Briefly, a pairwise comparison between all subpopulations was conducted with 1000 permutations. Significant ligand-receptor interaction pairs were generated and relevant ones were manually selected for presentation.

### Gene ontology and pathway enrichment analysis

The significant expressed genes in each subpopulation or group were subjected to gene ontology analysis (http://geneontology.org/) for overrepresentation enrichment test by the PANTHER ™. Significant terms (FDR < 0.05) were selected out.

## ACKNWLEDGEMENTS

We are most grateful to all the women who agreed to donate their tissue samples for this study. We acknowledge Ms. Joyce Yuen the project nurse and all gynecologists at Queen Mary Hospital for the collection of the samples. We are also grateful to the staffs at the Genomics Core, Centre for PanorOmic Sciences (CPOS), The University of Hong Kong, for their assistance in the single cell gem preparation and sequencing. This study was supported by funding from the General Research Fund of the Research Grants Council, Hong Kong (GRF 17158316), the National Natural Science Foundation of China-Swedish Research Council Collaboration Research Programme (NSFC-VR 3181101648), the Shenzhen Knowledge Innovation Programme of the Shenzhen Science and Technology Innovation Commission (20180264) and The University of Hong Kong - Shenzhen Hospital Scientific Research Training Plan (HKUSZH20192003).

## AUTHOR CONTRIBUTIONS

D.C., R.W.S.C., K.G.D. and W.S.B.Y. conceived and designed this study; D.C., and W.S.B.Y. performed data analysis and interpretation; D.C. wrote the manuscript; R.W.S.C. conducted experiments, acquired and analyzed the data; E.H.Y.N. recruited subjects and arranged sample collection from recruited subjects; R.W.S.C., E.H.Y.N and W.S.B.Y. supported by grants. All reviewed and approved the manuscript.

## DECLARATION OF INTERESTS

All authors declared no potential conflicts of interest.

## SUPPLEMENTARY INFORMATION

Supplementary Information includes thirteen tables and six figures.

