## Supplementary material for "Revealing Cellular Heterogeneity and *In Vitro* Differentiation Trajectory of Cultured Human Endometrial Mesenchymal-like Stem Cells Using Single-cell RNA Sequencing": Table S1-S5, S8-S10, S13, Figure S1-S6

**Table S1.** Sample information

| Sample ID | Age | Phase | Description | Reagent version in 10X Genomics platform |
|---|---|---|---|---|
| M1 | 31 | Menstrual | Endometrial aspirates | Chemistry v3 |
| M2 | 40 | | | Chemistry v2 |
| M3 | 32 | | | Chemistry v2 |
| S1 | 49 | Secretory | Full-thickness endometrium from hysterectomy | Chemistry v3 |
| S2 | 52 | | | Chemistry v3 |
| S3 | 51 | | | Chemistry v2 |
| S4 | 49 | | | Chemistry v2 |
| S3C | 51 | | 14-day Clonogenic progenies of S3 | Chemistry v2 |

**Table S2.** Summary statistics for sequencing and mapping data of 8 samples

| Sample ID | Number of cells | Mean reads per cell | Median genes per cell | Total genes detected | Median UMIs per cell | Total number of reads | Percent mapped reads |
|---|---|---|---|---|---|---|---|
| M1 | 4,761 | 77,241 | 2,680 | 21,287 | 11,108 | 3.68E+08 | 54.50% |
| M2 | 1,661 | 298,758 | 5,738 | 20,489 | 35,493 | 4.96E+08 | 64.70% |
| M3 | 1,900 | 295,731 | 6,047 | 20,215 | 44,949 | 5.62E+08 | 66.90% |
| S1 | 7,149 | 67,603 | 5,106 | 21,839 | 25,244 | 4.83E+08 | 65.90% |
| S2 | 7,346 | 64,024 | 5,225 | 21,730 | 27,331 | 4.70E+08 | 71.30% |
| S3 | 1,826 | 255,035 | 5,156 | 19,727 | 29,754 | 4.66E+08 | 70.30% |
| S4 | 1,623 | 298,757 | 4,076 | 19,500 | 18,058 | 4.85E+08 | 70.20% |
| S3C | 3,172 | 145,576 | 4,769 | 20,714 | 31,273 | 4.62E+08 | 72.90% |

**Table S3**. Summary of the cell and gene filtering process of 7 eMSC samples

| **Procedure** | **Count** |
|---|---|
| Cells removed by total number of UMIs (3MAD) | 632 |
| Cells removed by number of detected genes (3MAD) | 1296 |
| Cells removed by reads mapped to mitochondrial genes (3MAD) | 3054 |
| Cells removed by reads mapped to ribosomal genes (3MAD) | 638 |
| Genes removed by number of expressed cells (< 1% cells) | 19985 |
| Genes removed by their ribosomal and mitochondrial identity | 147 |
| Remaining total cells post filtering | 20646 |
| Remaining genes post filtering | 13406 |

**Table S4**. Cell number distribution across subpopulations in each sample

| Subpopulation | M1 | M2 | M3 | S1 | S2 | S3 | S4 | Total cells |
|---|---|---|---|---|---|---|---|---|
| **SP1** | 27 | 217 | 580 | 3,656 | 2,242 | 260 | 335 | 7,317 |
| **SP2** | 10 | 656 | 276 | 675 | 2,159 | 20 | 37 | 3,833 |
| **SP3** | 1219 | 37 | 100 | 1,036 | 464 | 706 | 149 | 3,711 |
| **SP4** | 46 | 580 | 515 | 947 | 1,112 | 29 | 212 | 3,441 |
| **SP5** | 6 | 2 | 63 | 110 | 66 | 54 | 39 | 340 |
| **SP6** | 835 | 51 | 58 | 52 | 224 | 51 | 362 | 1,633 |
| **SP7** | 0 | 0 | 0 | 0 | 308 | 4 | 2 | 314 |
| **SP8** | 57 | 0 | 0 | 0 | 0 | 0 | 0 | 57 |

**Table S5**. Genes used to calculate scores for different categories

| **Categories** | **Genes** |
|---|---|
| Housekeeping | *GAPDH, RN18S1, ACTB, TBP, UBC* |
| Stem | *FGF2, LIF, SOX2, TERT, GDF5* |
| MSC | *CTNNB1, EGF, HGF, ICAM1, IFNG, IGF1, IL10, IL1B, IL6, ITGB1, KITLG, MMP2, NES, NUDT6, PTPRC, SLC17A5, TNF, VEGFA, VIM, VWF, JAG1, NOTCH1, GDF15, SMAD4, HIC1, TWIST2, ANPEP, CASP3, CD44, ENG, ERBB2, FUT4, FZD9, ITGA6, ITGAV, MCAM, NGFR, NT5E, PDGFRB, PROM1, THY1, VCAM1, BDNF, CD200, COL10A1, FGF18, DLX2, DLX5, PDGFRA* |
| Adipogenic | *PPARG, RHOA, CEBPA, CEBPB, LEPR* |
| Chondrogenic | *HAT1, ITGAX, KAT2B, SOX9, ACAN, COL2A1, MMP13, CSPG4, BMP4, TGFB1, TGFB3, BMP6, KDR, MSX1, MSX2, BMP2, PRRX1* |
| Osteognic | *ALPL, IBSP, SP7, BGLAP, BMP7, COL1A1, FGF10, HDAC1, PTK2, SMURF1, SMURF2, TBX5, RUNX2, FGF9, BMP4, TGFB1, TGFB3, BMP6, KDR, MSX1, MSX2, BMP2, PRRX1* |
| Neurogenic | *NES, GFAP, PAX6, SOX2, NCAM1, NEUROD1, VIM* |
| Immunosuppresion | *CCL2, CSF3, VEGFA, IL7* |

**Table S6**. DEGs identified in each subpopulation of eMSCs
Check the separate excel file **Table S6.**

**Table S7**. Gene ontology results of DEGs of each subpopulation of eMSCs
Check the separate excel file **Table S7.**

**Table S8**. Summary of the cell and gene filtering process of paired sample (S3-S3C)

| **Procedure** | **Count** |
|---|---|
| Cells removed by total number of UMIs (3MAD) | 43 |
| Cells removed by number of detected genes (3MAD) | 820 |
| Cells removed by reads mapped to mitochondrial genes (3MAD) | 207 |
| Cells removed by reads mapped to ribosomal genes (3MAD) | 0 |
| Genes removed by number of expressed cells (< 1% cells) | 19567 |
| Genes removed by their ribosomal and mitochondrial identity | 4 |
| Remaining total cells post filtering | 3928 |
| Remaining genes post filtering | 13967 |

**Table S9**. Cell number distribution across subpopulations from S3-S3C

| Cluster | S3 | S3C | Total cells |
|---|---|---|---|
| **0** | 1 | 1,012 | 1,013 |
| **1** | 358 | 538 | 896 |
| **2** | 64 | 606 | 670 |
| **3** | 657 | 1 | 658 |
| **4** | 21 | 342 | 363 |
| **5** | 2 | 212 | 214 |
| **6** | 43 | 71 | 114 |

**Table S10**. Percentage of cells transitioning between clusters

| | **0** | **1** | **2** | **3** | **4** |
|---|---|---|---|---|---|
| **0** | - | 23.6 | 50.9 | 30.0 | 46.7 |
| **1** | 6.2 | - | 15.3 | 57.2 | 7.0 |
| **2** | 65.0 | 63.2 | - | 4.3 | 49.7 |
| **3** | 5.2 | 30.6 | 7.6 | - | 15.8 |
| **4** | 13.9 | 18.0 | 14.7 | 99.4 | - |

**Table S11**. Ligand-receptor interactions identified between different eMSC subpopulations
Check the separate excel file **Table S11.**

**Table S12**. Top 50 CD genes and HVGs in eMSCs, ADMSCs and WJMSCs
Check the separate excel file **Table S12.**

**Table S13**. Cell number distribution across subpopulations from 7 samples

| | **SP1 (7317)** | **SP2 (3833)** | **SP3 (3711)** | **SP4 (3441)** | **SP5 (340)** | **Total (18642)** | |
|---|---|---|---|---|---|---|---|
| ***LEPR*** | 890 | 450 | 1107 | 496 | 56 | 2999 | 16% |
| ***NES*** | 5329 | 2546 | 2875 | 2738 | 209 | 13697 | 73% |
| ***CSPG4*** | 2631 | 806 | 1180 | 1509 | 127 | 6253 | 34% |
| ***PDGFRA*** | 4922 | 2282 | 3201 | 2273 | 194 | 12872 | 69% |
| ***CD24*** | 378 | 213 | 537 | 235 | 30 | 1393 | 7% |
| ***DPP4*** | 353 | 224 | 401 | 209 | 69 | 1256 | 7% |
| ***ICAM1*** | 1347 | 576 | 1539 | 678 | 278 | 4418 | 24% |
| ***SUSD2*** | 136 | 46 | 114 | 73 | 15 | 384 | 2% |

*genes expressed with UMI > 0 in each cell were counted

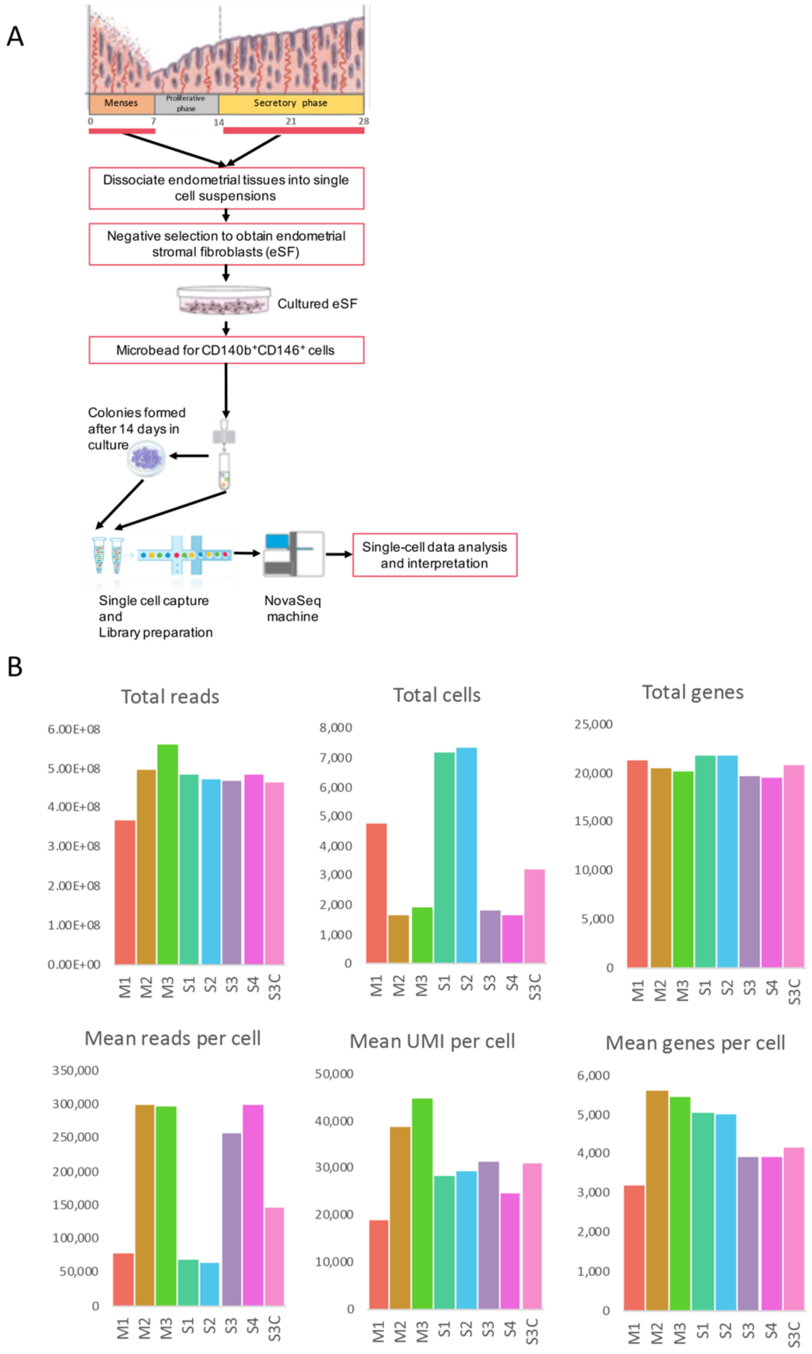


**Figure S1. Workflow and sequencing statistics.** (A) Workflow of this study; (B) Summary of sequencing statistics for seven samples is shown for six parameters.

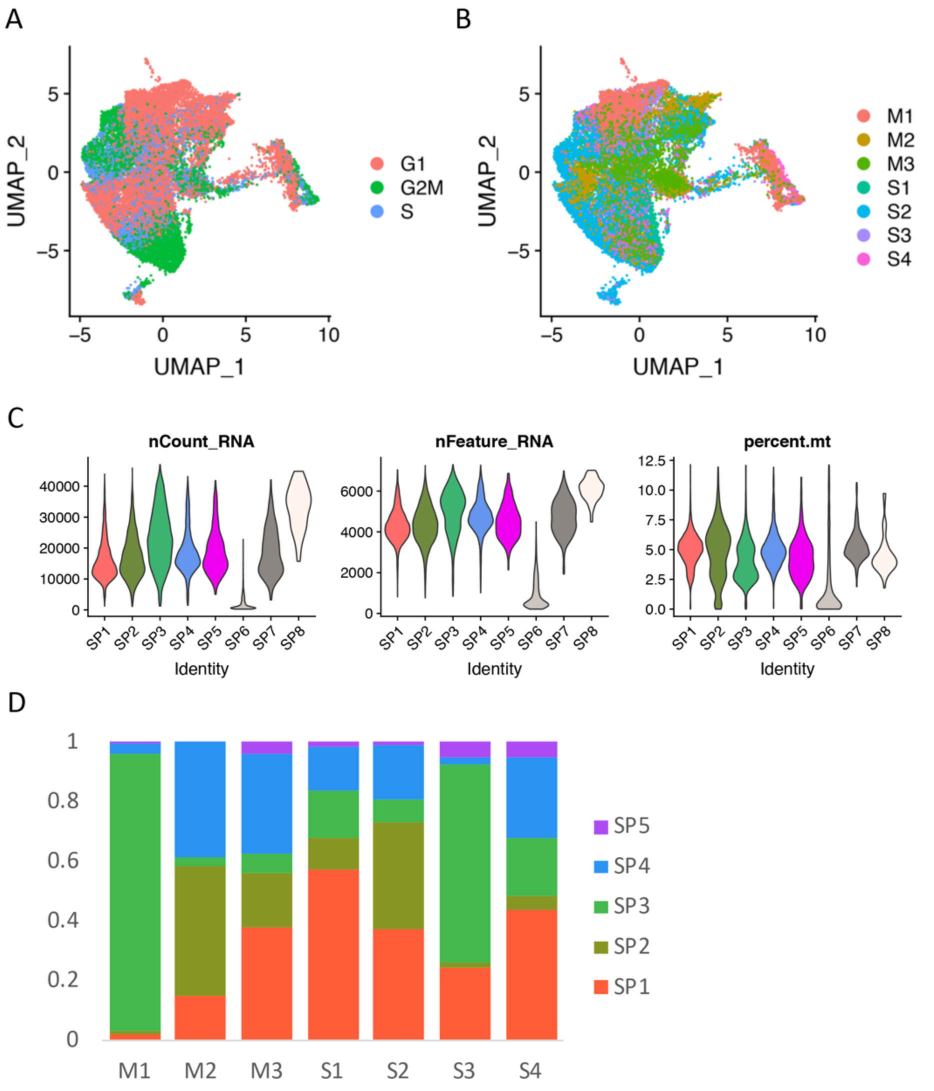


**Figure S2. Overview of primary clusters assigned for 7 eMSC samples.** (A-B) 2D UMAP plot of cells colored (A) by cell cycle phases and (B) by sample after correction. Each point represents one cell; (C) Violin plots for total UMIs per cell, total genes per cell and mitochondrial reads per cell in each subpopulation; (D) Cell number distribution across subpopulations in each sample.

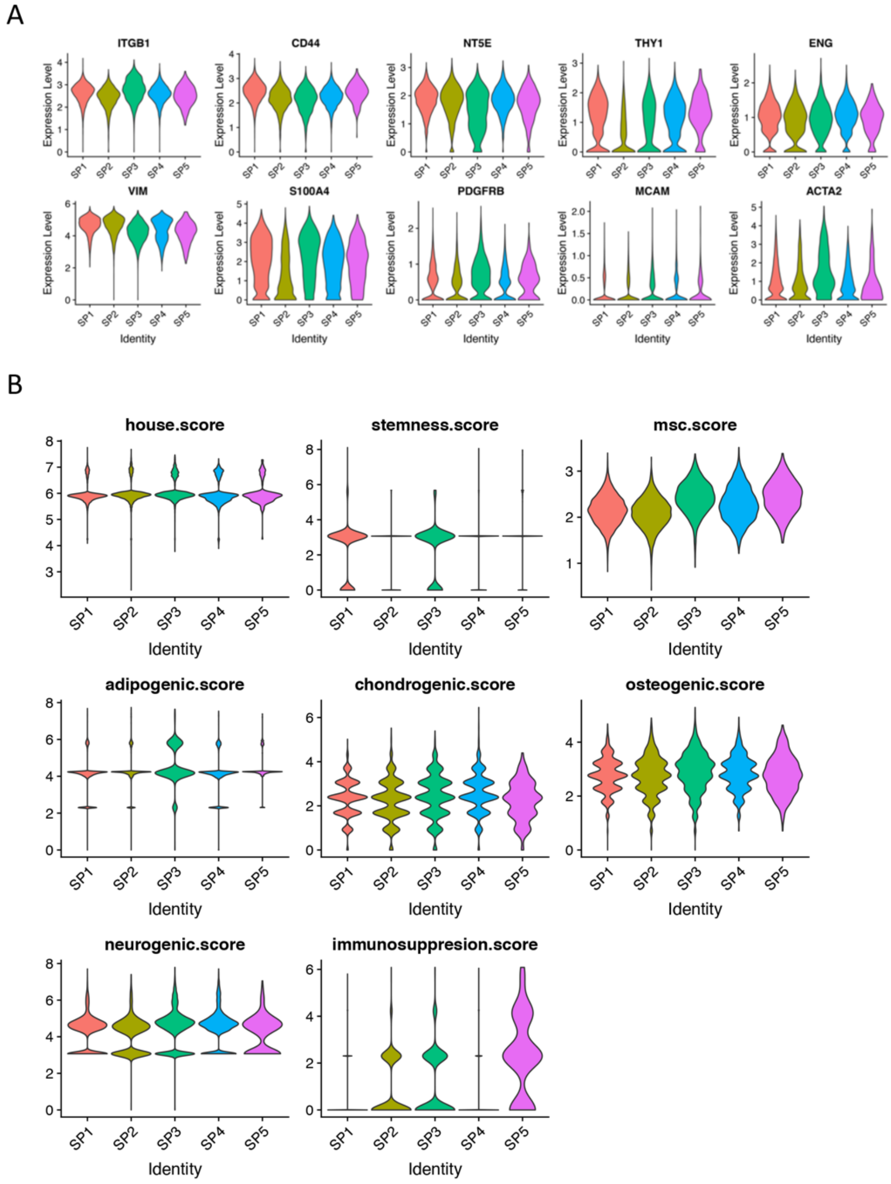


**Figure S3. Characterization of 5 candidate subpopulations for 7 eMSC samples.** (A) Violin plots for MSC markers, stromal cell markers and pericyte markers in candidate subpopulations; (B) Violin plots for different scores in candidate subpopulations. The score for each cell was obtained by averaging the expression of corresponding gene set. The genes used for calculating different scores was shown in Table S5.

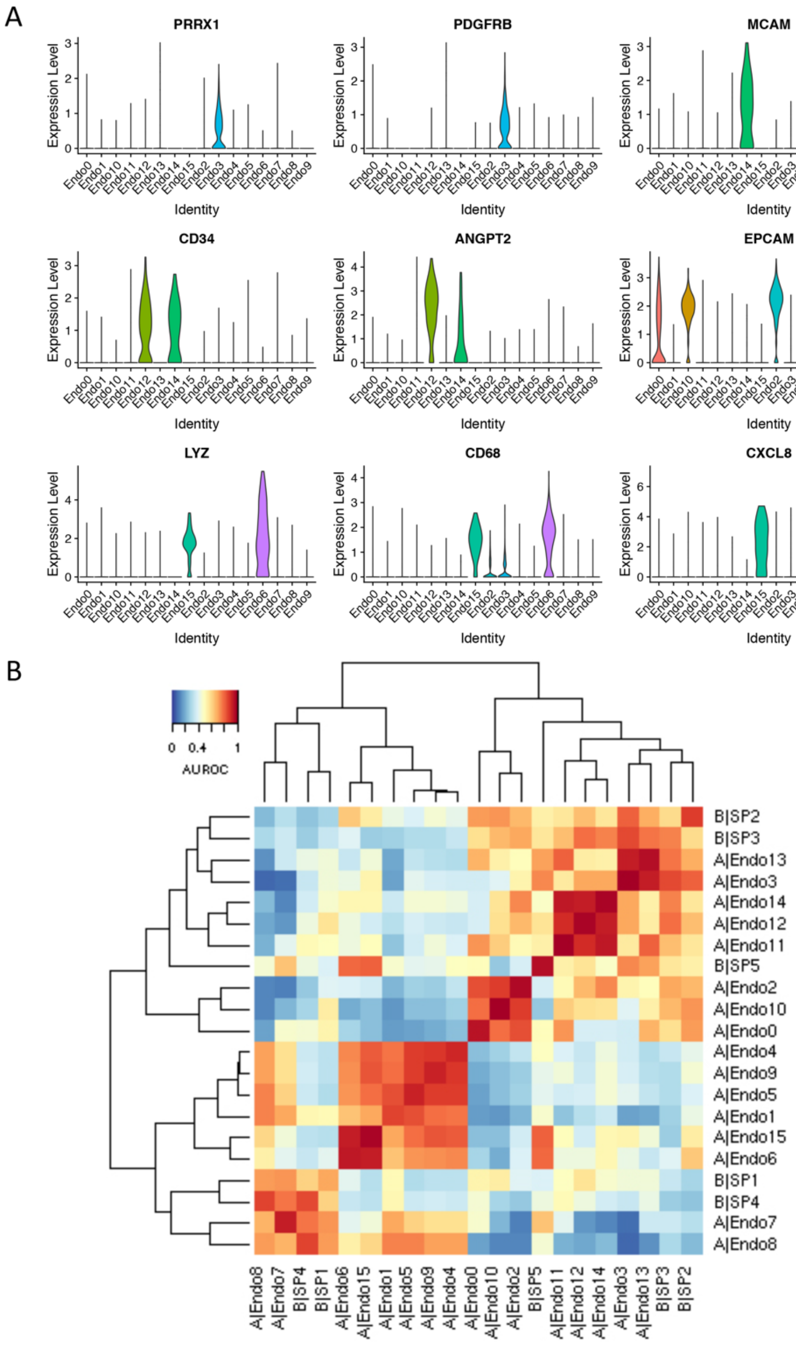


**Figure S4. Comparing eMSC subpopulations to primary bulk eMSC and primary endometrium.** (A) Violin plots for marker gene expression that mark stromal cells (PRRX1, PDGFRB), endothelial cells (CD34, ANGPT2, MCAM), epithelial cells (EPCAM), and macrophages (LYZ, CD68, CXCL8) in primary endometrium (In total, 16 subpopulations were identified from primary endometrium); (B) Heatmap of correlation on mean AUROC scores (from MetaNeighbor analysis) among cell subpopulations from primary endometrium (set A: Endo0-Endo15) and cultured eMSCs (set B: SP1-SP5). (AUROC scores greater than 0.7 indicates high linkage between the two subpopulations. AUROC scores of pairs: SP2-Endo3 = 0.81, SP3- Endo3 = 0.85, SP1-Endo7 = 0.73, SP4-Endo8 = 0.86, SP5-Endo15/SP5-Endo6 = 0.81.

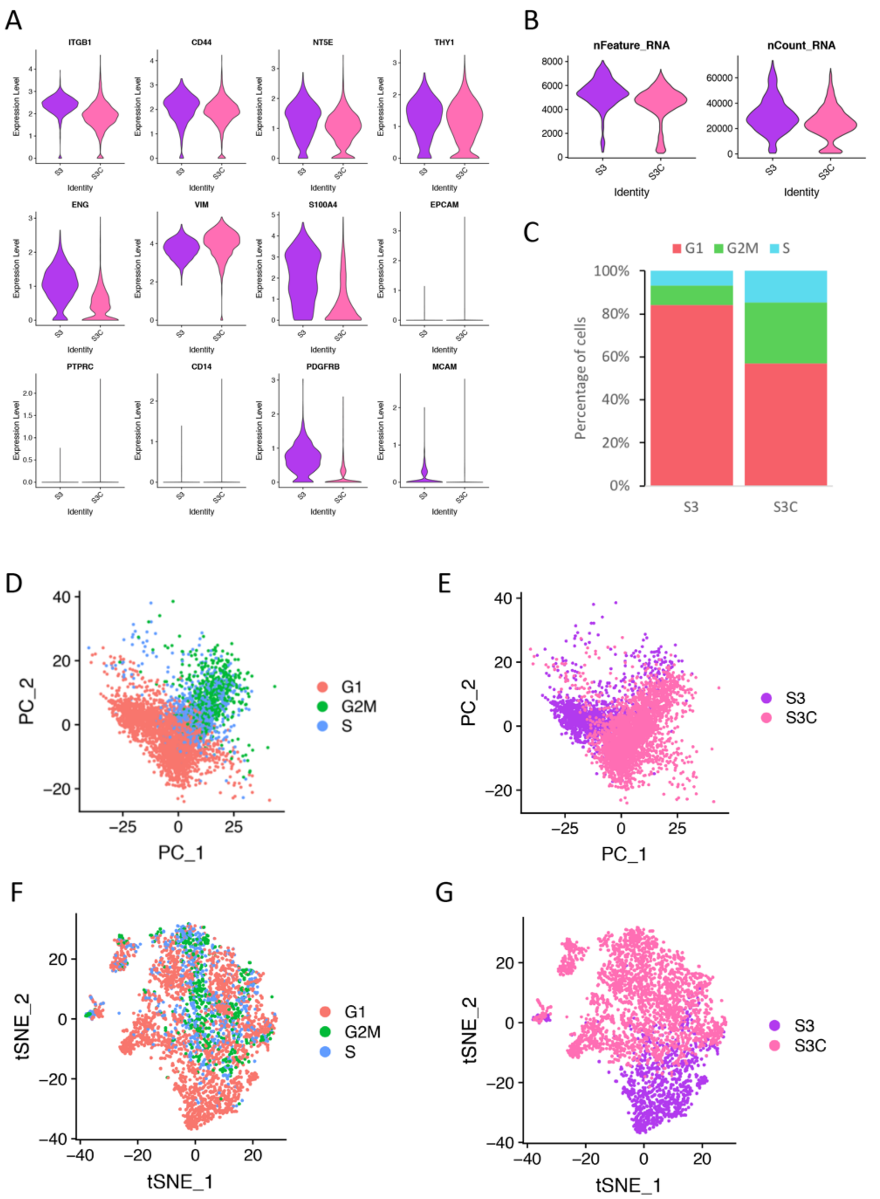


**Figure S5. Characterization and clustering of single cells from eMSC sample S3 and its clonogenic sample S3C.** (A) Violin plots for MSC markers, stromal cell markers, epithelial cell marker, immune cell marker, endothelial cell marker and eMSC markers in samples S3 and S3C; (B) Violin plots for total UMIs per cell and total genes per cell in each sample; (C) Phases of cell cycle distribution in samples S3 and S3C; (D, E) Cell distribution at PC1 and PC2 colored (C) by cell cycle effects and (D) by sample before correction; (F, G) tSNE plots of cells colored (E) by cell cycle phases and (F) by sample after correction.

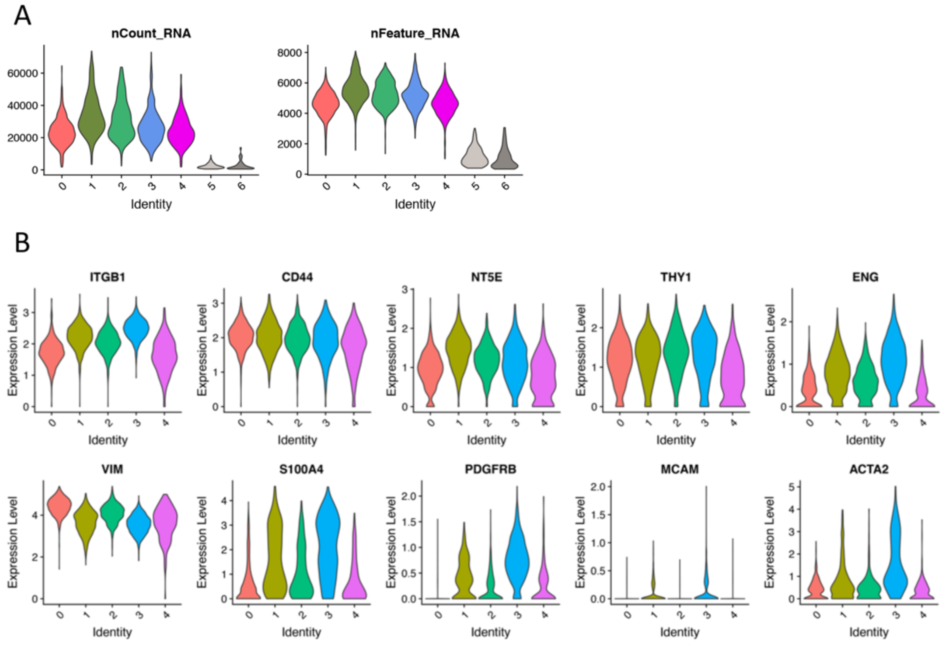


**Figure S6. Characterization of clusters assigned for S3 and S3C.** (A) Violin plots for total UMIs per cell and total genes per cell in primary clusters; (B) Violin plots for MSC markers, stromal cell markers, and pericyte markers in final clusters identified from S3 and S3C.
